# Phenotypic screening using synthetic CRISPR gRNAs reveals pro-regenerative genes in spinal cord injury

**DOI:** 10.1101/2020.04.03.023119

**Authors:** Marcus Keatinge, Themistoklis M. Tsarouchas, Tahimina Munir, Juan Larraz, Davide Gianni, Hui-Hsin Tsai, Catherina G. Becker, David A. Lyons, Thomas Becker

## Abstract

Acute CRISPR/Cas9 targeting offers the opportunity for scalable phenotypic genetic screening in zebrafish. However, the unpredictable efficiency of CRISPR gRNA (CrRNA) activity is a limiting factor. Here we describe how to resolve this by prescreening CrRNAs for high activity *in vivo*, using a simple standardised assay based on restriction fragment length polymorphism analysis (RFLP). We targeted 350 genomic sites with synthetic RNA Oligo guide RNAs (sCrRNAs) in zebrafish embryos and found that almost half exhibited > 90% efficiency in our RFLP assay. Having the ability to preselect highly active sCrRNAs (haCRs), we carried out a focussed phenotypic screen of 30 macrophage-related genes in spinal cord regeneration and found 10 genes whose disruption impaired axonal regeneration. Four (*tgfb1a, tgfb3, tnfa, sparc*) out of 5 stable mutants subsequently analysed retained the acute haCR phenotype, validating the efficiency of this approach. Mechanistically, lack of *tgfb1a* leads to a prolonged immune response after injury, which inhibits regeneration. Our rapid and scalable screening approach has identified functional regulators of spinal cord regeneration, and can be applied to study any biological function of interest.

**HIGHLIGHTS:**

- Synthetic CRISPR gRNAs are highly active
- *in vivo* pre-screening allows rapid assessment of CRISPR gRNA activity
- Phenotypic CRISPR screen reveals crucial genes for spinal cord regeneration
- *tgfb1a* promotes spinal regeneration by controlling inflammation

## INTRODUCTION

Embryonic and larval zebrafish are well suited to phenotypic screens of complex biological phenomena in living vertebrates. These screens assess how chemical or genetic manipulations affect biological processes of interest (Lam and Peterson, 2019). Zebrafish have been extensively employed in random mutagenesis-based discovery screens, but the emergence of CRISPR/Cas9 technologies now allows for specific gene targeting (Hwang et al., 2013). CRISPR-based approaches now also allow scalable assessment of gene function in zebrafish, because phenotypes of interest can be observed already in acutely injected, mosaically mutated embryos (Jao et al., 2013; Kuil et al., 2019; Shah et al., 2015). This is, in principle, a highly efficient screening approach, because it avoids time-consuming breeding to homozygosity (at least two generations with a generation time of 2-3 months each) of stable mutants that may or may not have relevant phenotypes (Hu et al., 2019; Pei et al., 2018).

The wide range in efficiency of guide RNAs, however, increases the possibility of failure to detect phenotypes of interest due to inefficiency of guides (false negatives). Compensating for that by injecting multiple guides targeting the same gene carries the risk of increasing off-target effects (false-positives) (Shah et al., 2016; Tsai et al., 2015; Wu et al., 2018). Therefore, we reasoned that developing a pipeline that includes pre-screening of guide RNAs for high activity would allow us to inject fewer guide RNAs and only those that were highly active. This would reduce the potential for off-target, false positive effects, as well as for false negative findings resulting from inactivity of guide RNAs. For a pre-screening approach to be feasible, we considered it important to use guides likely to have high activity and to have a high degree of freedom in target site selection. Moreover, we wanted to develop a rapid read-out measure to test for guide efficiency, based on standardised PCR and restriction enzyme-based analyses, to facilitate pre-screening in a scalable manner.

We tested the suitability of synthetic guides for such an approach, which others have recently shown to be reliably active in zebrafish (Hoshijima et al., 2019). In a synthetic RNA Oligo system, a generic RNA Oligo TracrRNA forms a complex with a bespoke RNA Oligo CrRNA (herein referred to simply as sCrRNA). As opposed to conventional *in vitro* transcription, sCrRNA design does not need to take preferred sequences for RNA polymerases into account. However, the GG of the protospacer adjacent motif (PAM) is still required.

To validate our pre-selection screening approach, we targeted 30 genes implicated in macrophage function to test how they affected spinal cord regeneration in larval zebrafish. Previous work has shown that the presence of macrophages is crucial for axonal reconnection in the injured larval spinal cord and recovery from paralysis (Tsarouchas et al., 2018). Specifically, blood-derived macrophages control the injury site environment by reducing the number of anti-regenerative neutrophils and by mitigating the injury-induced expression of pro-inflammatory cytokines, such as *Il-1b* by neutrophils and other cell types. However, the mechanisms and signals by which macrophages keep neutrophils and *il-1b* expression in check during regeneration are largely unknown.

Here, through our screen, we verified 4 genes as positive regulators of successful spinal cord regeneration in larval zebrafish and found that *tgfb1a* is necessary to limit the pro-inflammatory phase of the immune reaction. In summary, we present a rapid sCrRNA design and pre-selection process (summarized in Box 1) that facilitates rapid and robust phenotypic screening in vivo.

### Box 1 Recommendations for phenotypic screen workflow using sCrRNAs

1. *Design 4 sCrRNA per gene of interest.*

- do not use prediction programmes for efficiency
- target 5’ and/or conserved regions
- design all sCrRNAs such that restriction enzyme recognition sites are targeted
2. Identify haCRs

- co-inject 4 sCrRNAs per gene to teste guide efficiency.
- Use RFLP to identify haCRs (>90% efficiency sCrRNAs). Re-iterate if necessary.
3. Perform phenotypic screen

- co-inject up to two haCRs per gene and determine phenotypic read-out.
4. Validate in mutants

- generate stable mutants with early stop codons and determine phenotypic read-out.

## RESULTS

We aimed to explore the versatility of sCrRNAs for phenotypic screening in zebrafish. sCrRNAs have been reported to be more active than conventionally in vitro transcribed gRNAs, but this is based on a limited number of observations (Hoshijima et al., 2019). Here we quantitatively explore the activity of 350 sCrRNAs *in vivo* and demonstrate the use of sCrRNAs of pre-determined high activity in a focused phenotypic screen in spinal cord regeneration.

### sCrRNA activity is variable in vivo

We targeted 350 genomic sites (supplemental data file 1) in genes related to macrophage/microglia function and neurodegeneration and determined their activity towards the targeted sites by injecting into the zebrafish zygote. To do this efficiently, we targeted recognition sites for restriction enzymes, such that we could use resistance to enzyme digestion in restriction fragment length polymorphism analysis (RFLP) as a proxy for mutagenesis activity at the target site (Fig. 1A). Activity was determined by measuring band intensities of undigested and digested band and calculating their ratio. This ratio, from 0 (no activity) to 1 (complete target mutagenisation) was expressed as percentage efficiency. For example, a ratio of 0.9 was thus expressed as 90% efficiency. We defined as a highly active sCrRNA (haCR) those with an efficiency of > 90%. We favoured recognition sites for *bsl1, xcm1* and *bstxi* restriction enzymes, because these enzymes contain the necessary PAM sites for CrRNAs in their recognition sequences. Moreover, the recognition sites of these enzymes are also relatively large (>10 nucelotides), consisting of mostly redundant sequences. This makes them sensitive to formation of insertion and deletion mutations (indels) but also resistant to potential single nucleotide polymorphisms.

**Figure 1.**
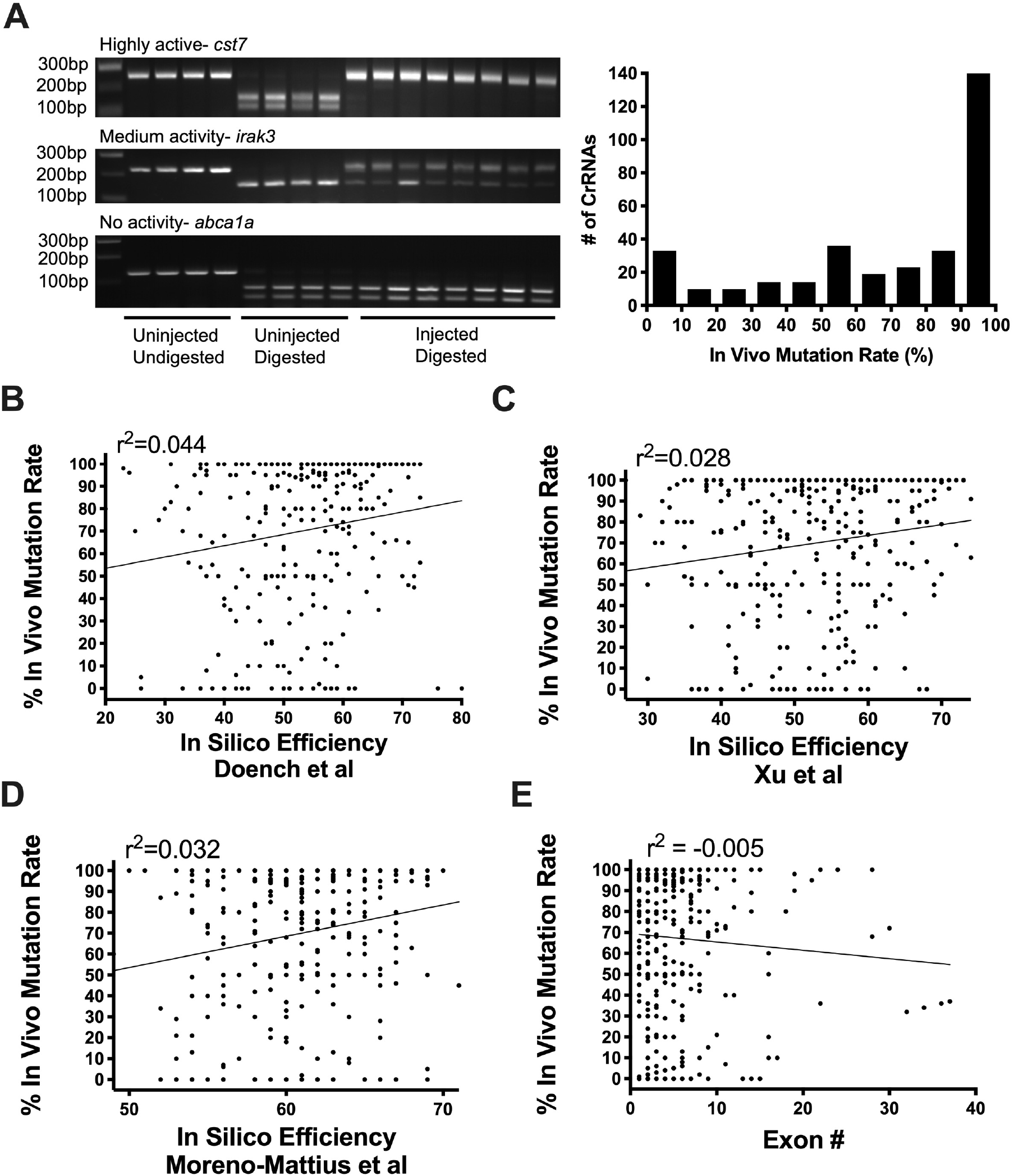
Many sCrRNAs are highly active. **A:** Example gels to assess sCrRNA in vivo mutation rate by resistance to restriction enzyme digest (RFLP) are shown. These indicate > 90% mutation rate (top), medium mutation rate (middle) and no detectable mutation rate (bottom). Each lane is derived from one animal. The chart shows the distribution of > 350 sCrRNAs by in vivo mutation rate. **B-D:** Activities of individual sCrRNAs show weak correlation with predicted in silico efficiency using different prediction rules (see results). **E:** sCrRNA activities are not correlated with their relative position to the start codon *in vivo*.

To increase the likelihood that CrRNAs were function-disrupting, we preferentially targeted the 5’ end of genes, increasing the chance that frameshift-inducing indels would lead to early stop codons likely to severely disrupt protein translation. In cases in which suitable 5’ target sites were not available, we targeted established functional domains. The subsequent RFLP analysis indicated that 44% of all tested sCrRNA were haCRs (Fig. 1A). Nonetheless, the activity of sCrRNA is variable, with many of them showing low activity (106 of 350 sCrRNAs showed < 50% activity), indicating that pre-screening guide RNAs ahead of phenotypic analyses is essential.

We next wanted to determine whether higher rates of haCR detection could be achieved by application of prediction rules for in vitro transcribed CrRNAs. Using CHOP-CHOP (Labun et al., 2019), we tested prediction guidelines which assume that GC content, single and dinucleotide identity at each position improves efficiency (Doench et al., 2016), that cytosine at the −3 position, adenines at −5/-12 and guanines at −14/-17 are favourable for high activity (Xu et al., 2015), and that guanine enrichment and adenine depletion increases targeting efficiency (Moreno-Mateos et al., 2015). Correlation between predictions and actual efficiencies observed in our RFLP-based assays were weak for all of these (R^2^-values between 0.028 - 0.044; Fig. 1B-D). Our preference for the 5’ end of the gene did also not bias sCrRNA for high activity (Fig. 1E). Hence, conventional design rules do not increase already high haCR detection rates, with the activity of individual sCrRNAs remaining largely unpredictable. This further indicates the benefits of a rapid robust pre-screening of activity of guides designed with less stringent criteria.

### haCRs efficiently ablate gene function

For phenotypic screening it is important that levels of functional protein are strongly reduced. This may best be achieved by frameshift mutations that can statistically be assumed for almost 66% of all alleles (1 - 1/3) for injection of one haCR (Moreno-Mateos et al., 2015). We aimed to use 2 haCRs per gene, where available, because the total proportion of frameshifts can be estimated to be 89% [1- (1/3 * 1/3)] per allele, so 79% for bi-allelic frameshifts in a cell (0.89 * 0.89). In addition, also in-frame indels may not be tolerated, particularly when known functional domains are targeted. Designing and injecting more than two haCRs could increase the bi-allelic frameshift rate (to 92% of cells for three haCRs), while increasing the possibility of off-target actions.

We used direct sequencing to test the prediction of substantial frameshift activity by using two haCRs for a test gene (Fig. S1A-C). The gene was chosen such that the two haCRs were in close proximity to each other (Cas9 cut sites 79bp apart) to be able to determine frameshifts in the same sequencing analysis. haCR#1 produced frameshifts in 24 of 33 (72%) clones and haCR#2 in 20 of 33 (60%). When both sites were taken into account simultaneously, the frameshift rate was increased to 87% of the alleles (29/33) (Fig. S1A-C), which is close to the theoretical value of 89% for a single allele.

To test whether haCR targeting would produce a substantial reduction in protein expression over both alleles, we targeted *hexb*, which codes for the lysosomal enzyme and microglial marker hexosaminidase, with two haCRs (Hickman et al., 2013), because a quantifiable enzyme assay was available to us. haCR#1 reduced enzyme activity by 63% and haCR#2 by 80%. Combining the haCRs reduced enzyme activity by 80%, not more than haCR#1 alone (Fig. S1D). Hence, one haCR can already lead to strongly reduced protein function, but using two haCRs increases the likelihood of consistent and efficient gene disruption.

### Phenotypic screening

To determine the usefulness of screening with sCrRNAs of pre-determined high activity, we decided to target macrophage function in spinal cord regeneration in a highly focused screen using haCRs. We have previously shown that macrophages are pivotal to spinal cord regeneration by controlling the inflammatory response to injury (Tsarouchas et al., 2018). Therefore, we targeted 30 macrophage-related genes (Table S1), for which we had identified at least 1 haCR in a functional domain or 2 haCRs.

We have previously shown that alterations of spinal cord regeneration can be effectively assessed in a simple read-out by determining the percentage of larvae with continuity of axonal labelling across the spinal injury site (called “bridging”) at 24 and 48 hours post-lesion (hpl) after a lesion at 3 days post-fertilisation (dpf) (Fig. 2A). This rapid way of quantification correlates with more complex measurements, such as the thickness of the axon bridge, and with the degree of recovery of swimming function animals show after injury (Tsarouchas et al., 2018). Here, for our haCR screen, we used animals with transgenically labelled axons (Xla.Tubb:DsRed) to visualise spinal cord bridging without the need to perform additional labelling (Fig. 2A). For duplicated genes (*slc2a5, timp2* and *lpl*), both paralogs were targeted simultaneously to avoid potential compensation by upregulation of the remaining paralog (El-Brolosy et al., 2019).

Of the 30 targeted genes, we found 10 ‘hits’ that significantly reduced proportions of larvae with axon bridging at 48 hpl (Fig. 2B). One of these (*tnfa*), described in a previous study (Tsarouchas et al., 2018), served as a positive control. The hit rate for genes targeted with one haCR (4 hits of 8 genes) in essential functional domains was not lower than that for genes targeted by two haCRs (6 hits of 22 genes), indicating that one haCR can be sufficient to produce phenotypes (see discussion).

**Figure 2.**
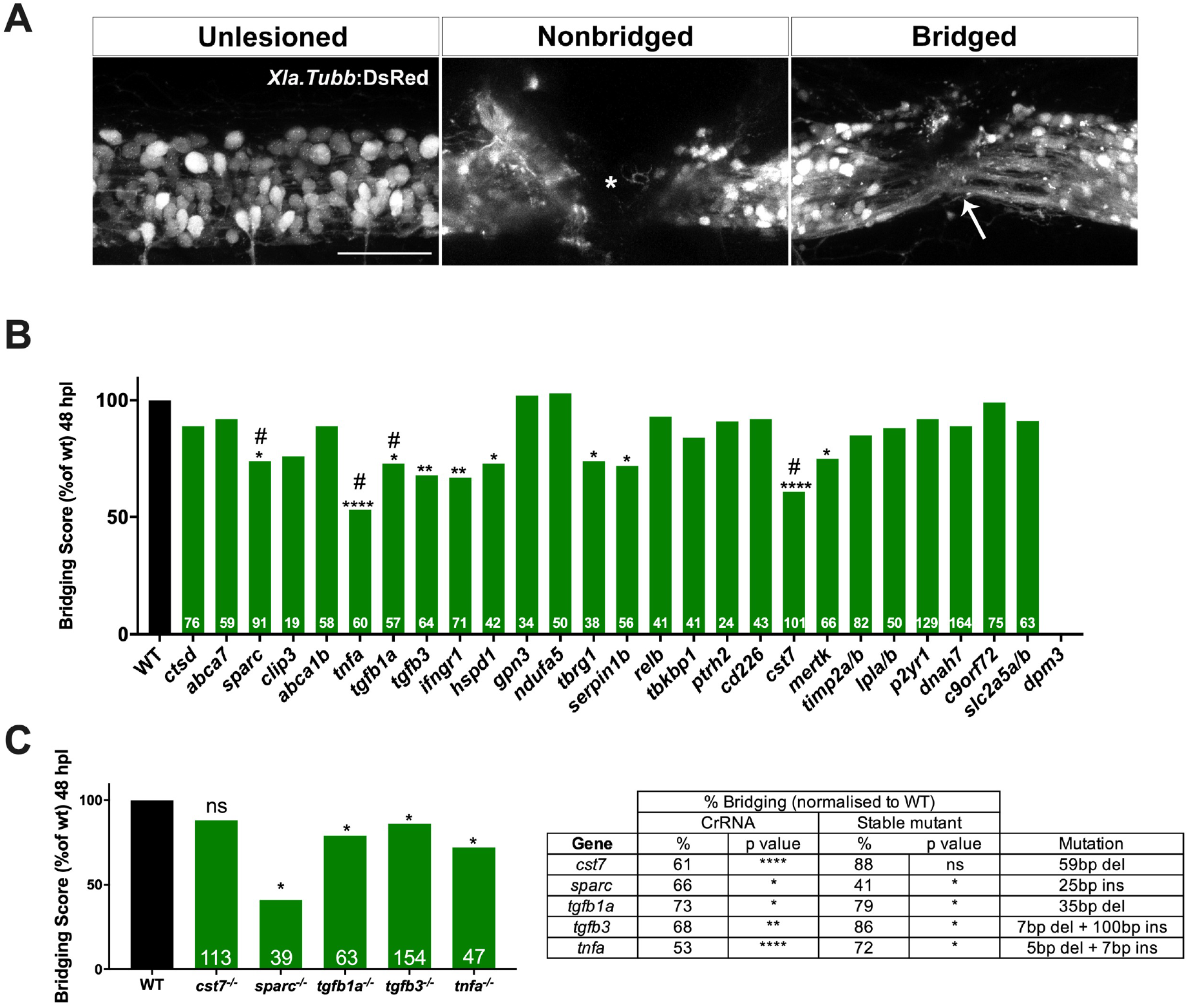
Phenotypic screening reveals modifiers of spinal cord regeneration. **A:** Example images of unlesioned, non-bridged (star indicates gap of neuronal labeling) and bridged spinal cord (white arrow) are shown (lateral views). Scale bar = 50 μm. **B:** Results of spinal cord regeneration screen for all screened genes at 48 hpl are shown. Significant reductions in bridging, normalised to control lesioned animals, are observed for *cst7* (p < 0.0001), *sparc* (p = 0.04), *tgfb1a* (p = 0.03), *tgfb3* (p = 0.005), *tnfa* (p < 0.0001), *ifngr1* (p = 0.0013,) *hspd1* (p = 0.011), *tbrg1* (p = 0.0494, *serpinb1* (p = 0.0279), and *mertk* (p = 0.0195); * indicates significance at 48 hpl; # indicates significance at 24 hpl (see Fig. S2); number of larvae per experiment are indicated at the bottom of each bar. For *dpm3* no viable larvae could be raised. A single sCrRNA targeting a key functional domain was used to target *ctsd, abca7, sparc, clip3, abca1b, tnfa, tgfb1a* and *tgfb3*. Two sCrRNAs were used to target all remaining genes. **C:** Mutant analysis confirms phenotypes for *sparc* (24 hpl, p = 0.024), *tgfb1a* (48 hpl, p = 0.019), *tgfb3* (48 hpl, p = 0.043) and *tnfa* (48 hpl, p = 0.024), but not for *cst7* (48 hpl, p = 0.079). The table compares the magnitude of effects between acute injection and in mutants. Fischer’s exact test was used for all analyses.

### Validation of hit genes

To validate hit genes, we raised stable mutants from haCR-injected embryos for select genes, *tgfb1a, tnfa, sparc*, and *cst7*. These genes were chosen, because we observed reduced rates of larvae with bridged injury sites at the early and late time points of analysis after spinal injury for these (Fig. S2). This indicates that these genes may be essential for regeneration from an early time point. We added *tgfb3* to the list for mutant validation, despite only showing reduced bridging at 48 hpl, because of the possibly related functions *of tgfb3* and *tgfb1a* as anti-inflammatory cytokines.

All mutations produced premature stop codons, confirmed by direct sequencing, and therefore are likely to abrogate gene function (Fig. S3). For phenotypic analysis, we outcrossed mutants to wildtype animals and analysed incrosses of these heterozygous animals (F3 generation) to mitigate the risk of carrying forward background mutations. Proportions of larvae with bridged lesion sites were assessed against wildtype siblings.

With the exception of *cst7*, all stable mutants showed impaired axon bridging of the spinal lesion site at 48 hpl (Fig. 2C). *tgfb1a* mutants showed comparable magnitudes of the bridging phenotype to acute haCR injection (mutant: 79% larvae with bridged injury site compared to wildtype; acute injection: 73%), whereas *sparc* mutants were more strongly affected than acutely injected larvae (Mutant: 41% larvae with bridged injury site compared to wildtype; acute injection: 66%). Mutants for *tgfb3* (86% larvae with bridged injury site compared to wildtype; acute injection: 68%) and *tnfa* (72% larvae with bridged injury site compared to wildtype; acute injection: 53%) showed a milder phenotype than acutely haCR injected larvae (Fig. 2C). Despite some quantitative differences between acute injections and stable mutants, we confirmed impaired spinal cord regeneration for 80% of the tested hit genes also in mutants.

### tgfb1 controls post-injury inflammation

We decided to analyse the consequences of targeting *tgfb1a* and *tgfb3* on spinal cord regeneration in more detail, because both are expressed in macrophages in the injury site (Tsarouchas et al., 2018) and may be needed to mitigate the lesion-induced inflammation. Both neutrophil numbers and *il-1b* expression levels are elevated in the macrophage-less *irf8* mutant at 48 hpl (Tsarouchas et al., 2018). Therefore, we assessed numbers of neutrophils, macrophages and levels of *il-1b* expression after *tgfb1a* and *tgfb3* haCR injection at this timepoint.

To visualise macrophages in the injury site, we used the *mpeg1:GFP* transgenic line. We did not find any changes in macrophage numbers after injection of haCRs to *tgfb1a* or *tgfb3* in lesioned animals, compared to lesioned controls (Fig. 3A,B), indicating that these genes are not necessary for macrophage to populate the injury site.

**Figure 3.**
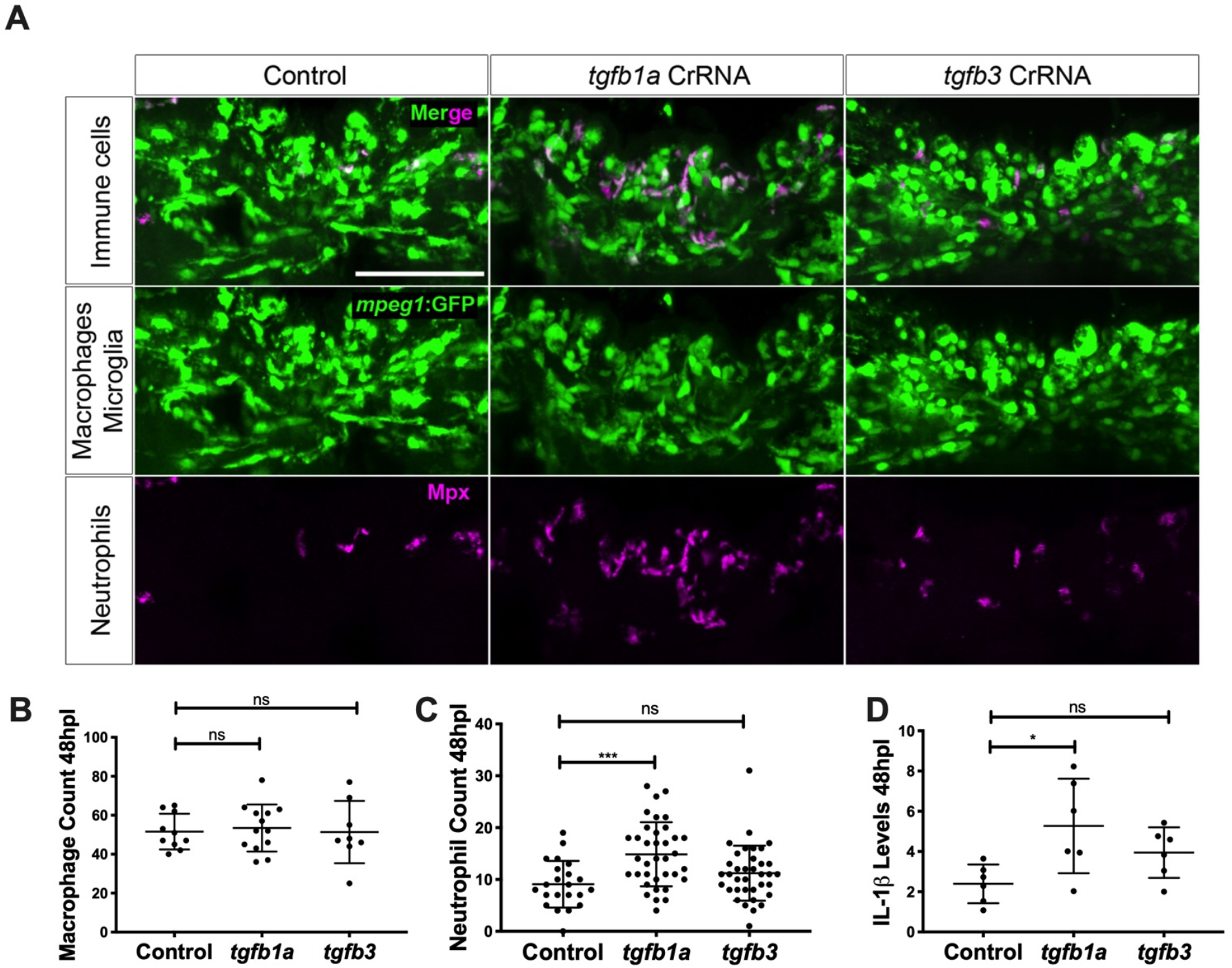
Loss of *tgfb1a* leads to prolonged inflammation. **A:** Lateral views of lesion sites in larval zebrafish are shown with the indicated markers and experimental conditions at 48 hpl. **B-C:** Quantifications show that numbers of macrophages were not altered by injecting any of the indicated haCRs (B; one-way ANOVA with Bonferoni’s multiple comparison test), neutrophils were increased in number in *tgf1b* haCRs injected animals (one-way ANOVA with Bonferroni’s multiple comparison test, p = 0.0006), but not in *tgfb3* haCRs injected animals (p = 0.32). **D:** Animals injected with *tgfb1a* haCRs, but not those injected with *tgfb3* haCRs (p = 0.36), displayed marked increases in *il1b* expression levels in the lesion site compared to lesioned controls at 48 hpl (one-way ANOVA with Bonferroni’s multiple comparison test, P = 0.0211). Error bars represent standard error of the mean (SEM). Scale bar in A = 50 μm. All transcript levels were normalized to uninjected, unlesioned controls.

To visualise neutrophils, we used an antibody to Mpx in the same animals as above. Injection of *tgfb1a* haCRs led to an increase of 63% in neutrophil numbers after injury compared to lesioned controls (Fig. 3A,C). Lesioned animals injected with *tgfb3* haCRs showed a trend towards an increase in neutrophil counts of 23% compared to lesioned controls, but this did not reach significance (p > 0.05; Fig. 3A,C).

To measure levels of *il1b* expression, we used qRT-PCR on the isolated injured trunk region (Tsarouchas et al., 2018). This showed a 120% increase in *il1b* expression at 48 hpl after injection of *tgfb1a* CrRNA, but no effect for *tgfb3* (Fig. 3A,D), compared to control lesioned animals. The continued presence of high numbers of neutrophils and high *il1b* expression at 48 hpl after *tgfb1a* haCR targeting indicates prolonged inflammation, shown to inhibit regeneration (Tsarouchas et al., 2018).

In summary, our haCR screen has found several macrophage-related genes to be involved in successful spinal cord regeneration in zebrafish and reveals *tgfb1a* and *tgfb3* as promoting spinal cord regeneration, at least in part, by controlling neutrophil numbers and *il1b* expression.

## DISCUSSION

We show that pre-screening sCrRNAs for activity is an effective way of eliminating the inherent variability of CrRNA activity from phenotypic screens in zebrafish. In a screen of genes associated with macrophage function, we demonstrate roles of *tgfb1a, tgfb3, tnfa*, and *sparc* for optimal spinal cord regeneration in zebrafish mutants.

### Pre-screening for activity improves phenotypic screening with sCrRNAs

To observe phenotypes after acute sCrRNA injection, it is crucial to significantly reduce protein expression. We calculated that two haCRs per gene should result in a strong disruption of gene function and we confirmed this by direct sequencing and assessing protein activity. Moreover, phenotypes can also be found after injection of only one haCR targeting essential functional domains. Knowing that sCrRNAs used are highly active strongly reduces the risk of false negative results. Moreover, using maximally two sCrRNAs reduces the potential for off-target effects associated with higher numbers of injected CrRNAs in screening efforts using CrRNAs of unknown efficiency (Tsai et al., 2015). Although we found phenotypes in our pilot screen using only 1 haCR for certain genes and efficiency of one haCR can be very high, we argue that if 2 haCRs can be identified per gene, this is more likely to be effective, as the relative effect on protein coding function is likely to be consistently higher. This is especially the case for poorly characterised genes, for which the consequences of in frame mutations are difficult to predict. Furthermore, certain cell types may exhibit homeostatic responses such that unaffected cells could outcompete cells with essential functions disrupted by gene targeting.

Pre-screening has become feasible due to the availability of synthetic CrRNAs, which have a higher likelihood of being highly active than in vitro transcribed sgRNAs. Indeed, our rate of haCRs of 44% is substantially higher than the 1-10% reported for in vitro transcribed sgRNAs (Gagnon et al., 2014; Moreno-Mateos et al., 2015). sCrRNA targeting is also superior to using in vitro transcribed sgRNAs, because the targeted sequence does not need to contain initiation sites for transcription enzymes that are needed for conventional in vitro transcription. This allows for a larger freedom in target selection.

As a reason for a large proportion of sCrRNAs being highly active, it has been shown that an exact match to the target sequence without any addition of bases, introduced by in vitro transcription, is important (Gagnon et al., 2014; Hoshijima et al., 2019). Moreover, as a two-part system, consisting of sCrRNA and TracrRNA, sCrRNAs may have different Cas9 and DNA binding activities than sgRNAs.

However, despite the high rate of haCRs, activity of individual sCrRNAs is highly variable and difficult to predict. We applied current prediction rules for CrRNA activity to our collection of sCrRNAs and found only weak correlations. Hence, prescreening is essential to eliminate variability in targeting efficiency.

Phenotypic screening of regeneration with haCRs is limited to genes with no essential developmental function. We observed excessive mortality when targeting *dmt3*, probably because of essential developmental roles of the gene (Marchese et al., 2016). It is likely that we did not observe more instances of non-developing embryos, because we targeted mostly macrophage-related genes and macrophages are not essential for early development (Shiau et al., 2015).

The genes for our screen were chosen because of their relation to the function of macrophages, which play a crucial role in controlling the inflammation in successful spinal cord regeneration in zebrafish (Tsarouchas et al., 2018). We were therefore not surprised to find a high hit rate of 33%. We confirmed a role for the genes in spinal cord regeneration in stable mutants for 4 out of 5 genes (80%). This indicates that the screening paradigm has a relatively low rate of false positive findings.

Theoretically, our assay could also detect genes that are inhibitors of regeneration, by showing accelerated axonal bridging after haCR injection. Indeed, stimulation of the immune response with bacterial lipopolysaccarides accelerates axonal regeneration, indicated by increased percentages of larvae with a bridged injury site at 24 hpl (Tsarouchas et al., 2018). However, we did not find any of the haCR targeting of 30 genes to lead to an increase in bridging efficiency at that time point (Fig. S2). Hence, none of the targeted genes are inhibitors of regeneration.

Lack of confirmation of a phenotype in the *cst7* mutant could be due to off-target effects of the sCrRNAs on other genes after acute injection. To safeguard against such effects, CrRNAs with minimal predicted alternative binding sites should be given priority for screening. Even on-target activity can lead to deletion of large DNA regions, leading to unintended effects (Thomas et al., 2019). However, it is also possible that mutants are hypomorphs or produce dominant proteins of altered activity. This may also explain quantitative differences in the bridging phenotype between acutely sCrRNA injected animals and stable mutants.

### The immune reaction is essential for spinal cord regeneration

The phenotypes of *tgfb1a* sCrRNA-injected animals resembled that of a lack of macrophages (Tsarouchas et al., 2018). In the macrophage-less *irf8* mutant, neutrophils show a slower clearance rate from the injury site after the peak at 2 hours post-lesion, such that their number is increased relative to wildtype animals at 48 hpl. Likewise, levels of *il-1b*, expressed mainly by neutrophils, peak at 4-6 hours after injury and rapidly decline thereafter, but do so more slowly in the *irf8* mutant. As *tgfb1a* and *tgfb3* are typical anti-inflammatory cytokines (Li et al., 2017) and are expressed by macrophages in zebrafish (Tsarouchas et al., 2018) and mammals (McTigue et al., 2000) after spinal injury, Tgfb signalling may be a part of the mechanism by which macrophages control the inflammation and thereby promote spinal cord regeneration. However, *tgfb1a* and *tgfb3* are also expressed by other cell types (Tsarouchas et al., 2018) and may interact with cell types other than neutrophils, such as astrocytes (Rathore et al., 2012) or the neurons directly (Li et al., 2017; Vidal et al., 2013).

Tnfa is mainly produced by macrophages in the spinal injury site and has been shown to be necessary for larval spinal cord regeneration using pharmacological inhibition of Tnfa release and acute sCrRNA injection in zebrafish (Tsarouchas et al., 2018). We confirm these findings here in a stable *tnfa* mutant.

Sparc is produced by macrophages and organises the collagen matrix (Bradshaw, 2016). Expression of *Sparc* by transplanted olfactory ensheathing glia in mammalian spinal cord injury has been shown to be beneficial to regeneration (Au et al., 2007). In zebrafish larvae, regenerating axons grow in close contact with fibrils of extracellular matrix containing growth-promoting ColXII (Wehner et al., 2017). Hence Sparc protein may contribute to this axon growth promoting matrix in zebrafish.

Mutation of *cst7* did not impair spinal cord regeneration. The gene codes for Cystatin F, which is an inhibitor of cathepsin C, a lysosomal protease (Hamilton et al., 2008). Hence, activity of *cst7* may be related to phagocytosis of debris in macrophages. Interestingly, we have shown previously that the phagocytic function of macrophages is not critical for axonal bridging (Tsarouchas et al., 2018), which may in part explain the lack of an observe phenotype in the *cst7* mutant.

In conclusion, we propose to improve phenotypic screening in zebrafish with CRISPR/Cas9 by using superior synthetic RNA oligo CrRNAs and by pre-screening these for high activity before performing a phenotypic screen (Box 1). We demonstrate this approach by finding immune system related genes that are essential for successful spinal cord regeneration.

## MATERIAL AND METHODS

### Animal husbandry

All zebrafish lines were kept and raised under standard conditions and all experiments were approved by the British Home Office (project license no.: 70/8805).

The following lines were used: WIK wild type zebrafish (Johnson and Zon, 1999), Tg(Xla.Tubb:DsRed)^zf14826^, abbreviated as Xla.Tubb:DsRed (Peri and Nusslein-Volhard, 2008); Tg(*mpeg1*:EGFP)^gl22^, abbreviated as *mpeg1:GFP* (Ellett et al., 2011).

### Crispr/Cas9 design and injection

All CrRNAs were designed so a restriction enzyme recognition sequence overlapped the Cas9 cut site. CrRNAs and TracrRNA were purchased from Merck KGaA (Germany, Darmstadt). Cas9 (M0386M, NEB, Ipswich USA) and all restriction enzymes were purchased from NEB (Ipswich USA). Cas9 was diluted to 7 μM with diluent buffer B (NEB, Ipswich USA) on arrival and stored at −20 °C. RNA oligos were re-suspended to 20 μM with nuclease free water, and stored at −20 until use. For in vivo testing of CrRNA activity, 1 nl of an injection mixture composed of 1 μl of each CrRNA (up to 4), 1 μl TracrRNA, 1 μl Cas9 and 1 μl Fast Green (Alfa Aesar, Heysham, UK), was injected into the yolk of single cell stage embryos.

### Generation of stable mutants

CrRNAs targeting exon 1 were injected into the yolk of one-cell stage embryos. The CrRNA target sites for *tgfb1a* and *tgfb3* were 5’ ATGGCTAAAGAGCCTGAATCCGG and 5’GAATCCATCCAGCAGATCCCTGG, respectively. *tnfa* was targeted with 5’ ACAAAATAAATGCCATCATCGGG, *sparc* with 5’ CTAAACCATCACTGCAAGAAGGG and *cst7* with two sCrRNAs, 5’TTCTGCAGAGCTCCTGGGATCGG and 5’ TAAAAGAGTAAGTTCCAGTCAGG, to generate a larger deletion. Founders were identified and out crossed to WT (F1), then crossed to WT again to generate the F2 generation. F2 heterozygous individuals were crossed to Xla.Tubb:DsRed or wildtype to generate the F3 generation. All spinal cord lesion assays were performed on an F3 heterozygous incross.

*The tgfb1a* line was genotyped with primers F 5’ GATTTGGAGGTGGTGAGGAA and R 5’ TCGCTCAGTTCAACAGTGCTAT. The *tgfb3* line was genotyped with primers F 5’ GGGTCAGATCCTCAGCAAAC and R 5’ GAGATCCCTGGATCATGTTGA, the *sparc* line with F 5’ TGCCTAAACCATCACTGCAA and R 5’ ATGCTCGAAGTCTCCGATTG, and the *cst7* line with F 5’ TTGTGTGCTCTTTGCTGTCTG and R’ CTGCACCTGTCTCTTTGCAC. The *tnfa* line was genotyped with primers F 5’ACCAGGCCTTTTCTTCAGGT and R 5’ AGCGGATTGCACTGAAAAGT followed by a *bstXI* digest.

### RFLP analysis

At 24 hpf, DNA from single embryos was extracted using 100 μl of 50 mM NaOH and 10 μl of Tris-HCl pH 8.0 as previously described (Wilkinson et al., 2013) All RFLP analyses were conducted on DNA of separate individuals and not pooled in order to accurately determine mutation rate.

PCR products were generated with BIOMIX red (BIOLINE, London, UK) and 1 μl of the respective restriction enzyme was added directly to the final PCR product for RFLP analysis, without the addition of extra buffers and incubated at the optimal temperature for each respective enzyme. 20 μl of digest were run on 2 % agarose gel (BIOLINE, London, UK) and imaged on a trans illuminator. Band intensities were calculated using imageJ (http://imagej.nih.gov/ij).

### Allele sequencing

PCR products from 8 injected embryos (24 hpf) were pooled and ligated into a StrataClone vector and transformed into competent cells, following the manufacturer’s instructions (Agilent, Santa Clara, USA). Positive colonies were identified and sequenced using M13 primers.

### Hexb activity assay

Hexb activity was determined as previously described (Keatinge et al., 2015). Briefly, 3 dpf embryos, 20 per group, were homogenised in 100 μl of nuclease free water. Each sample was diluted 1/10 with McIlvaine citrate-phosphate buffer pH 4.5 and activity assayed with 4-methylumbelliferyl-2-acetamido-2-deoxy-β-D-glucopyranoside (Sigma, Dorset, UK) in a plate reader.

### Immunohistochemistry on whole-mount larvae

All incubations were performed at room temperature unless stated otherwise. At the time point of interest, larvae were fixed in 4% PFA-PBS containing 1% DMSO at 4°C overnight. After washes in PBS, larvae were washed in PBTx. After permeabilization by incubation in PBS containing 2 mg/ml Collagenase (Sigma) for 25 min larvae were washed in PBTx. They were then incubated in blocking buffer for 2 h and incubated with primary antibody (anti-MPX, GeneTex, Irvine, California, USA) diluted in blocking buffer at 4°C overnight. On the following day, larvae were washed 3 x in PBTx, followed by incubation with secondary antibody diluted in blocking buffer at 4°C overnight. The next day, larvae were washed three times in PBTx and once in PBS for 15 min each, before mounting in 70% glycerol.

For whole mount immunostaining of acetylated tubulin to visualise the axons, larvae were fixed in 4% PFA for 1 h and then were dehydrated in 25% 50%, 75% MeOH in 0.1% Tween in PBS, transferred to 100% MeOH and then stored at 20°C overnight. The next day, head and tail were removed, and the samples were incubated in pre-chilled 100% acetone at −20°C for 10 min. Thereafter, larvae were washed and digested with Proteinase K (10 μg/ml) for 15 min at room temperature and re-fixed in 4 % PFA. After washes, the larvae were incubated with 4 % BSA in PBTx for 1 h. Subsequently, the larvae were incubated over two nights with primary antibody (acetylated tubulin). After washes and incubation with the secondary antibody, the samples were washed in PBS for 15 min each, before mounting in glycerol.

### Spinal cord injury

At 3 dpf, zebrafish larvae were anaesthetised in PBS containing 0.02 % aminobenzoic-acid-ethyl methyl-ester (MS222, Sigma), as described (Tsarouchas et al., 2018). Larvae were transferred to an agarose-coated petri dish. Following removal of excess water, the larvae were placed in a lateral position, and the tip of a sharp 30.5 G syringe needle (BD Microlance™) was used to inflict a stab injury or a dorsal incision on the dorsal part of the trunk at the level of the 15th myotome, leaving the notochord intact.

### Assessment of axonal phenotypes

Re-established axonal connections (“bridges”) were scored at the time point of interest in fixed immunolabelled samples and live transgenic animals. Larvae were directly visually evaluated using a fluorescent stereomicroscope (Leica M165 FC) or confocal imaging (Zeiss LSM 710, 880). Larvae were scored as described (Tsarouchas et al., 2018) with the observer blinded to the experimental condition. Briefly, a larva was scored as having a bridged lesion site when continuity of the axonal labeling between the rostral and caudal part of the spinal cord was observed. Continuity of labeling was defined as at least one fascicle being continuous between rostral and caudal spinal cord ends, irrespective of the fascicle thickness. Larvae in which the lesion site was obscured by melanocytes or the notochord was inadvertently injured were excluded from the analysis.

### Quantitative RT-PCR

Reverse transcription of 500 ng RNA was performed with the iSCRIPT kit (BIORAD, Hercules, USA). Standard RT-PCR was performed using 10 mM of each dNTP and each primer. qRT-PCR was performed at 58°C using Roche Light Cycler 96 and relative mRNA levels determined using the Roche Light Cycler 96 SW1 software. Samples were run in duplicates and expression levels were normalized to a β-actin control. Primers were designed to span an exon-exon junction using the Primer-BLAST (https://www.ncbi.nlm.nih.gov/tools/primer-blast/) software.

### Assessment of immune cell numbers

A volume of interest was defined centered on the lesion site from confocal images. The dimensions were: width = 200 μm, height = 75 μm (above the notochord), depth = 50 μm. Images were analysed using the Imaris (Bitplane, Belfast, UK) or the ImageJ software. The number of cells was quantified manually in 3D view, on at least three independent clutches of larvae, blinded to the experimental condition.

### Experimental design and statistical analysis

Image analysis was performed using ImageJ. Power analysis using G*Power (Faul et al. 2009), was used to calculate power (aim > 0.8) for the experiments and determine the group sizes accordingly. Statistical power was > 0.8 for all experiments. All quantitative data were tested for normality and analyzed with parametric and non-parametric tests as appropriate. The statistical analysis was performed using IBM SPSS Statistics 23.0. Shapiro-Wilk’s W-test was used in order to assess the normality of the data. Kruskal-Wallis test followed by Dunn’s multiple comparisons, One-way ANOVA followed by Bonferroni multiple comparisons test, two-way ANOVA, followed by Bonferroni multiple comparisons, t-test, Mann-Whitney U test or Fischer’s exact test were used, as indicated in the figure legends. *P< 0.05, **P< 0.01, ***P< 0.001, n.s. indicates no significance. Error bars indicate the standard error of the mean (SEM). The figures were prepared with Adobe Photoshop CC and Adobe Illustrator CC. Graphs were generated using GraphPad Prism 7.

## ACKNOWLEDGEMENTS

We thank Viviane Schulz for help with some experiments. This work was supported by a Wellcome Trust Senior Research Fellowship (102836/Z/13/Z) and by Biogen who provided funding via a scientific research agreement with DAL. Work in the Becker group is funded by the EU Cofund consortium NEURONICHE with contributions from MRC (Eranet Neuron Cofund), Spinal Research, and Wings for Life, as well as two project grants from the BBSRC (BB/R003742/1, BB/S001778/1).

## AUTHOR CONTRIBUTIONS

MK, TMT: Conceptualization, Data curation, Formal analysis, Validation, Investigation, Visualization, Methodology, Writing—original draft, Writing—review and editing; TM, JL: Investigation, Writing—review and editing; DG, HHT: Resources, Writing—review and editing; CGB, DL, TB: Conceptualization, Supervision, Funding acquisition, Investigation, Visualization, Writing—original draft, Project administration, Writing—review and editing.

## SUPPLEMENTARY DATA

### Supplemental tables

**Table S1:**
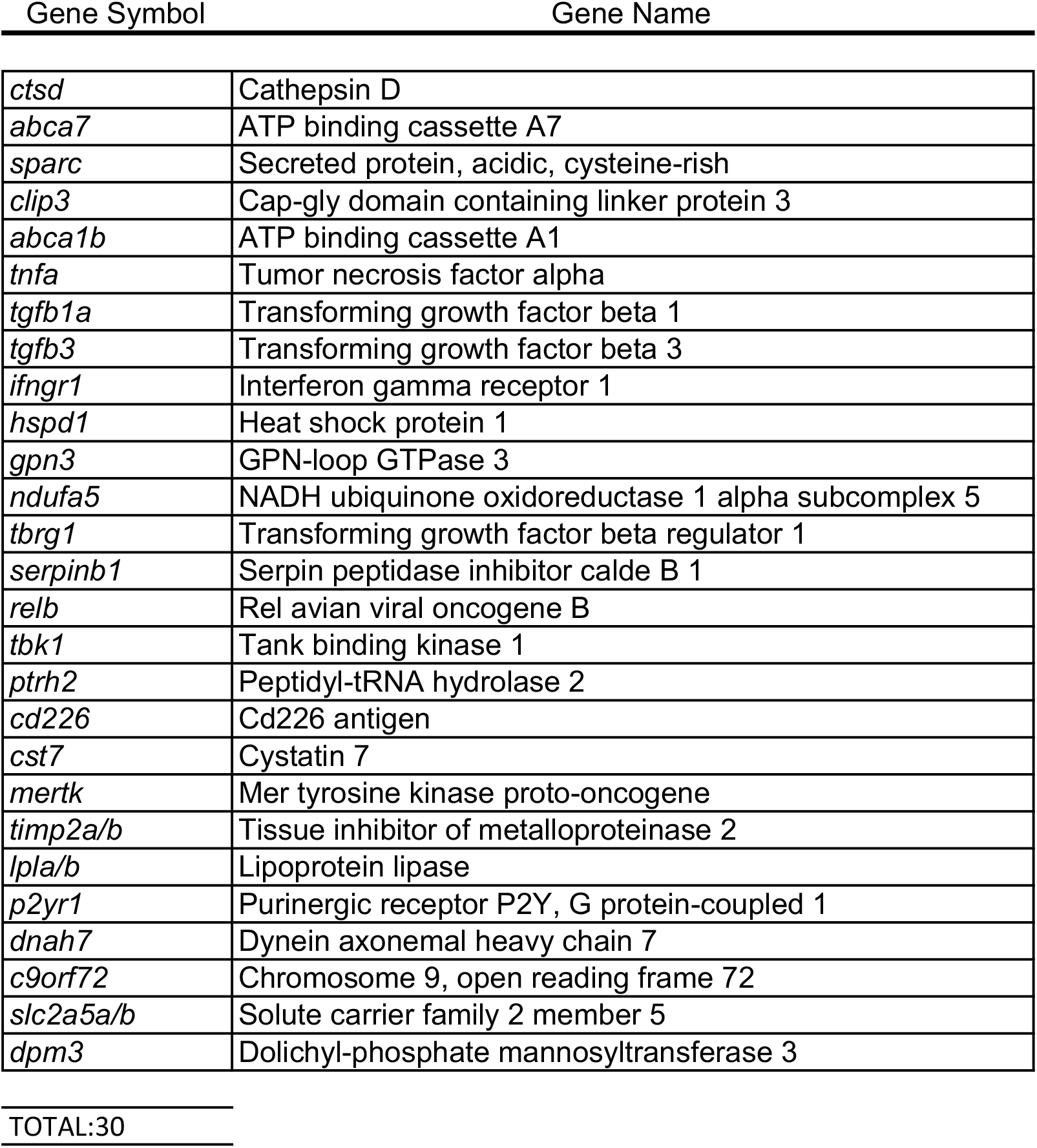
List of target genes for phenotypic screening in spinal cord repair. *slc2a5, timp2 and lpl* are duplicated in the zebrafish genome

| Gene Symbol | Gene Name |
| --- | --- |
| *ctsd* | Cathepsin D |
| *abca7* | ATP binding cassette A7 |
| *sparc* | Secreted protein, acidic, cysteine-rich |
| *clip3* | Cap-gly domain containing linker protein 3 |
| *abca1b* | ATP binding cassette A1 |
| *tnfa* | Tumor necrosis factor alpha |
| *tgfb1a* | Transforming growth factor beta 1 |
| *tgfb3* | Transforming growth factor beta 3 |
| *ifngr1* | Interferon gamma receptor 1 |
| *hspd1* | Heat shock protein 1 |
| *gpn3* | GPN-loop GTPase 3 |
| *ndufa5* | NADH ubiquinone oxidoreductase 1 alpha subcomplex 5 |
| *tbrg1* | Transforming growth factor beta regulator 1 |
| *serpinb1* | Serpin peptidase inhibitor calde B 1 |
| *relb* | Rel avian viral oncogene B |
| *tbk1* | Tank binding kinase 1 |
| *ptrh2* | Peptidyl-tRNA hydrolase 2 |
| *cd226* | Cd226 antigen |
| *cst7* | Cystatin 7 |
| *mertk* | Mer tyrosine kinase proto-oncogene |
| *timp2a/b* | Tissue inhibitor of metalloproteinase 2 |
| *lpla/b* | Lipoprotein lipase |
| *p2yr1* | Purinergic receptor P2Y, G protein-coupled 1 |
| *dnah7* | Dynein axonemal heavy chain 7 |
| *c9orf72* | Chromosome 9, open reading frame 72 |
| *slc2a5a/b* | Solute carrier family 2 member 5 |
| *dpm3* | Dolichyl-phosphate mannosyltransferase 3 |
| TOTAL:30 |  |

### Supplemental figures

**Fig. S1.**
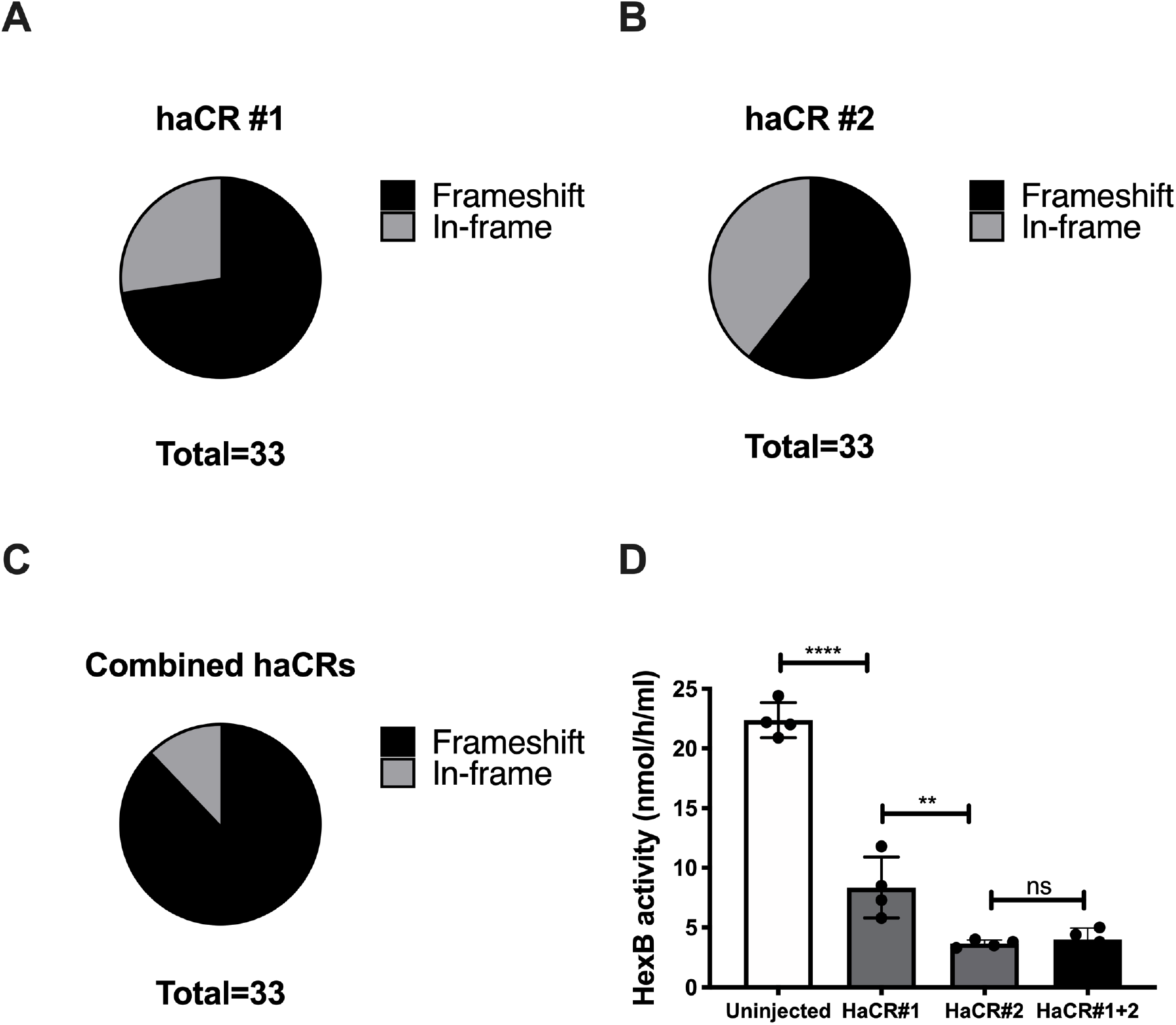
Injecting two haCRs simultaneously effectively disrupts gene function. **A-C:** Direct sequencing of mutant alleles in embryos injected with two haCRs per gene demonstrates induction of frameshift frequencies of 72% (A; haCr #1) and 60% (B; haCR #2), when either CrRNA is analysed individually. This rises to 87% when frame-shift frequencies are combined (C). **D:** At the protein level, dual haCR injection against *hexb* reduces enzyme activity by 80% *in vivo* (ANOVA with Tukey post-test; p < 0.0001). Single haCRs also reduce enzyme activity (p < 0.0001 for both). haCR#2 is more efficient than haCR#1 (p = 0.0051). Error bars represent SEM.

**Figure S2.**
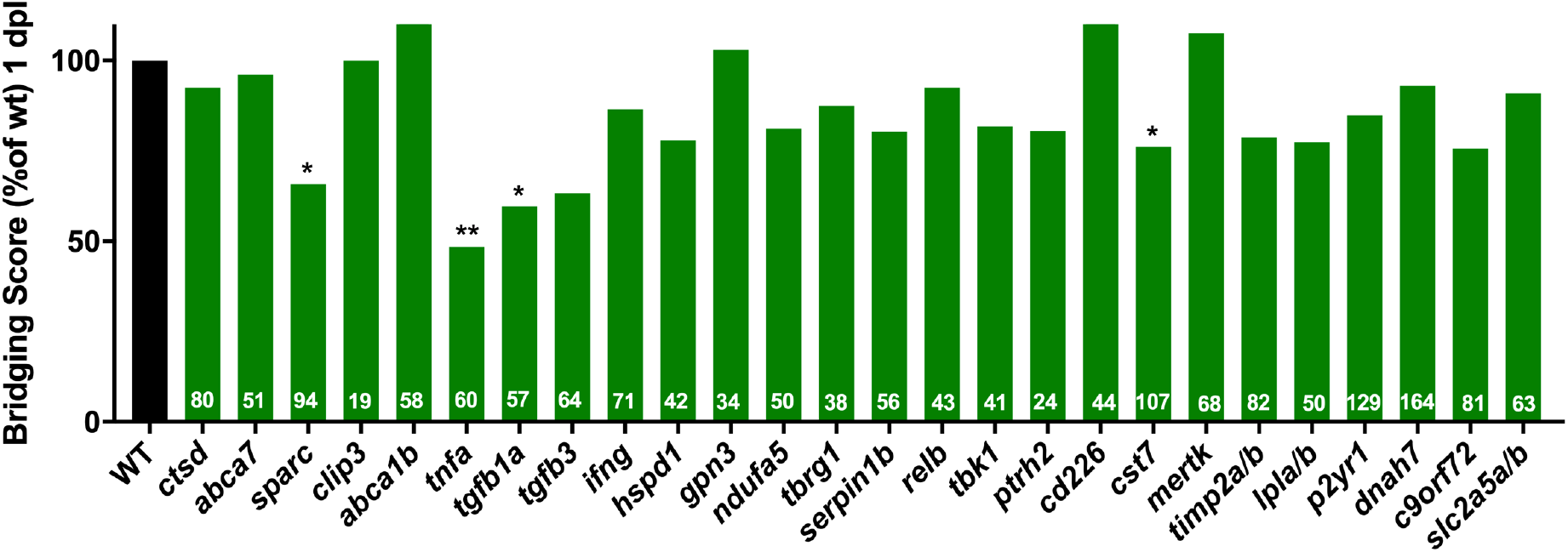
Significant inhibition of axon bridging is seen for some targeted genes at 24 hpl. Significant reductions in bridging were detected for acute injection of haCRs for *sparc* (p = 0.0424), *tnfa* (p = 0.0068), *tgfb1a* (p = 0.0225) and *cst7* (p = 0.0418) Fisher’s exact test was used for all analyses.

**Fig. S3.**
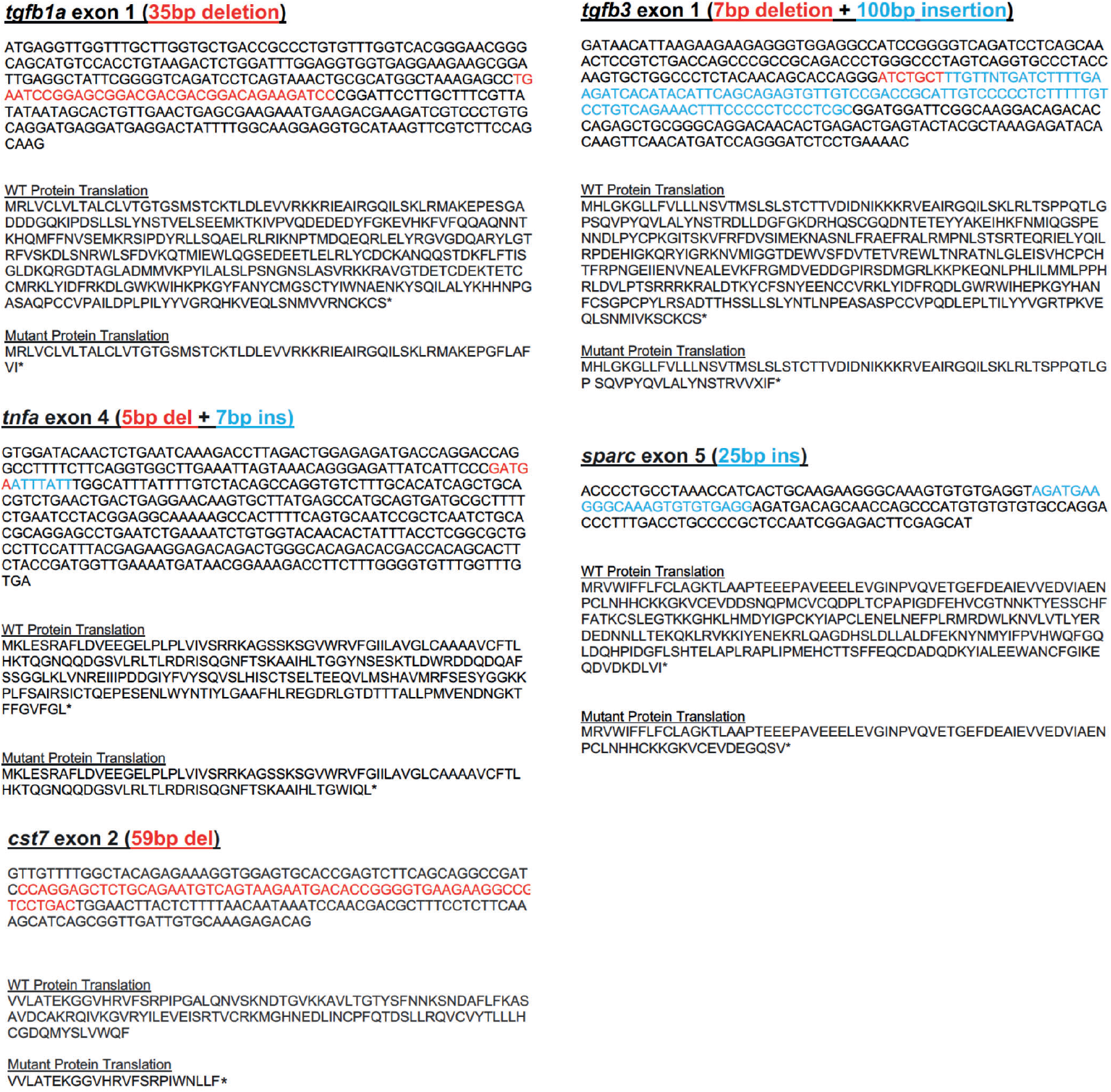
Mutations likely lead to non-functional protein. Deletions are shown in red and insertions are shown in blue. All stable mutations produce frameshifts (*cst7, tnfa, tgfb1a* and *sparc*) and premature stop codons, with the exception of the mutation in the *tgfb3* gene. The latter contains an in-frame indel in which the large quantity of inserted material contains a nonsense mutation.

